# A CRISPR-del-based pipeline for complete gene knockout in human diploid cells

**DOI:** 10.1101/2021.06.21.449335

**Authors:** Takuma Komori, Shoji Hata, Akira Mabuchi, Mariya Genova, Tomoki Harada, Masamitsu Fukuyama, Takumi Chinen, Daiju Kitagawa

## Abstract

The advance of CRISPR/Cas9 technology has enabled us easily to generate gene knockout cell lines by introducing insertion/deletion mutations (indels) at the target site via the error-prone non-homologous end joining repair system. Frameshift-promoting indels can disrupt gene functions by generation of a premature stop codon. However, there is growing evidence that targeted genes are not always knocked-out by the indel-based gene disruption. In this study, we established a pipeline of CRISPR-del, which induces a large chromosomal deletion by cutting two different target sites, to perform “complete” gene knockout efficiently in non-transformed human diploid RPE1 cells. By optimizing several procedures, the CRISPR-del pipeline allowed us to generate knockout cell lines harboring bi-allelic large chromosomal deletions in a high-throughput manner. Quantitative analyses show that the frequency of gene deletion with this approach is much higher than that of conventional CRISPR-del methods. The lengths of the deleted genomic regions demonstrated in this study are longer than those of 95% of the human protein-coding genes. Furthermore, the pipeline enables the generation of a model cell line having a bi-allelic cancer-associated chromosomal deletion. Overall, these data lead us to propose that the CRISPR-del pipeline is a high-throughput approach for performing “complete” gene knockout in RPE1 cells.

## Introduction

Gene knockout is a powerful technique for studying gene functions in experimental biology. Recent advances in genome editing technology have enabled us to conduct the gene knockout approach efficiently in a variety of organisms and cultured cells (Doudna and Charpentier, 2014; Hsu et al., 2014). Among the currently available genome editing tools, the CRISPR/Cas9 system, derived from the adaptive immune system of prokaryotes, is the most widely used tool for gene knockout. The CRISPR-associated protein Cas9 is an RNA-guided endonuclease that induces double-strand breaks (DSBs) at specific DNA loci (Jinek et al., 2012). These DSBs can be repaired via the efficient but error-prone non-homologous end joining (NHEJ) pathway, which frequently introduces small insertion/deletion mutations (indels) at the junction sites (Chang et al., 2017; Rouet et al., 1994). Frameshift-promoting indels generate premature termination codons (PTCs) downstream of the target site, leading to nonsense-mediated decay of the transcript or production of a nonfunctional truncated protein (Popp and Maquat, 2016).

Most of the widely applied approaches for CRISPR/Cas9-mediated gene disruption rely on the introduction of indels after DSB (Ran et al., 2013; Shalem et al., 2015), although there is accumulating evidence that this method would not always ensure a complete gene knockout. For example, Bub1, a spindle assembly checkpoint kinase, is known to be a “zombie” protein, which remains functional despite the presence of CRISPR/Cas9-generated indels (Meraldi, 2019; Rodriguez-Rodriguez et al., 2018; Zhang et al., 2019). Recent studies demonstrate that Bub1 “knockout” clones express alternatively spliced Bub1 mRNA (Rodriguez-Rodriguez et al., 2018) or low levels of an active Bub1 mutant harboring a small deletion (Zhang et al., 2019). Similarly, nonsense-associated altered splicing enables skipping of the exon harboring a PTC, thus allowing the production of functional proteins (Bagheri et al., 2022; Mou et al., 2017; Tuladhar et al., 2019). In addition to the altered splicing, alternative translation initiation is another often observed phenomenon in mutant clones generated by the indel-based method (Tuladhar et al., 2019). Several mechanisms, such as leaky scanning and internal ribosome entry, allow translation to start from an alternative initiation site on the edited transcript, in order to evade gene disruption caused by PTCs (Xu et al., 2020). Due to these cellular abilities to bypass PTCs, the indel-based method is not always the optimal strategy to achieve complete gene knockout.

On the other hand, the CRISPR-Cas9 deletion (CRISPR-del) is an alternative approach that can be applied to unambiguously disrupt a gene of interest (Canver et al., 2014; Raaijmakers and Medema, 2019; Xiao et al., 2013). CRISPR-del uses Cas9 and two different single guide RNAs (sgRNAs) to delete a large chromosomal region flanked by the two target sequences, thus ensuring a complete gene knockout. However, the efficiency of chromosomal deletions by CRISPR-del and the feasible deletion length in routine lab work have not been thoroughly investigated, especially in non-transformed human diploid cells.

In this study, we developed a pipeline of CRISPR-del for the use in high-throughput gene knockout in hTERT-immortalized RPE1 cell line, which is one of the most widely used human diploid cell lines. Two different quantitative analyses show that our CRISPR-del strategy is more efficient than previous CRISPR-del methods and is capable of deleting a very long region of genomic DNA in RPE1 cells. The length covers that of more than 95 % of human protein-coding genes. Furthermore, the CRISPR-del method can be used to create model human cell lines with cancer-associated large chromosomal deletions identified in patients. Taken together, this study leads us to propose that the CRISPR-del pipeline enables high-throughput complete gene knockout in non-transformed human diploid cells.

## Results

### An optimized CRISPR-del pipeline is a high-throughput approach for gene knockout in human diploid RPE1 cells

In order to apply CRISPR-del for high-throughput gene knockout in RPE1 cells, we first optimized several steps in the conventional method of CRISPR-del (Canver et al., 2014). To avoid the construction of sgRNA plasmids that can be time consuming, sgRNAs were synthesized via *in vitro* transcription from PCR-assembled DNA templates (Fig. 1a). We generally designed two different sgRNAs for each upstream and downstream target site flanking a deletion region, having 4 different combinations of sgRNA pairs. Each sgRNA pair was mixed with commercially available recombinant Cas9 protein, and the ribonucleoprotein (RNP) complexes were electroporated into RPE1 cells under an optimized condition. Delivery of CRISPR RNP via electroporation is known to be the most robust method in terms of editing efficiency and minimized off-target effects (Kim et al., 2014; Liang et al., 2015). After recovery from electroporation and genome editing, the deletion efficiency of each sgRNA pair was promptly analyzed by genomic PCR with primers designed to detect the expected deletion. From the cell pool of the sgRNA pair with the highest deletion efficiency, single cells were isolated into 96-well plates. The use of an automated single cell dispensing system based on piezo-acoustic technology resulted in a reliable clone isolation with high viability. After expansion for approximately 3 weeks in culture, each cell colony was detached from its well with Trypsin/EDTA solution, and split into two new 96-well plates. For one of them, the cells were cultured in normal medium and then subjected for genomic DNA extraction directly in the plate. In case of long-term storage of the other replicated plate, the detached cells in Trypsin/EDTA solution were directly mixed with three times volume of a DMSO-free cryopreservation medium, and the plate was placed and stored in a −80°C freezer. This procedure allows for genotyping analysis and cell expansion from the replicated plate at flexible timing. Two types of genotyping PCRs were conducted with the extracted DNA by using primers to detect either the WT or the deleted allele in a high-throughput manner for the whole plate. The PCR products were automatically analyzed by a microtip electrophoresis system to identify bi-allelic knockout clones showing an expected pattern of a deletion band and the absence of a WT band. Knockout clones were then expanded from the replicated plate and subjected to a second genomic PCR using their purified genomic DNA to confirm the expected deletion in both alleles. In summary, our optimizations in several procedures enable CRISPR-del to become a high-throughput method for generation of gene knockout cells.

**Fig.1:**
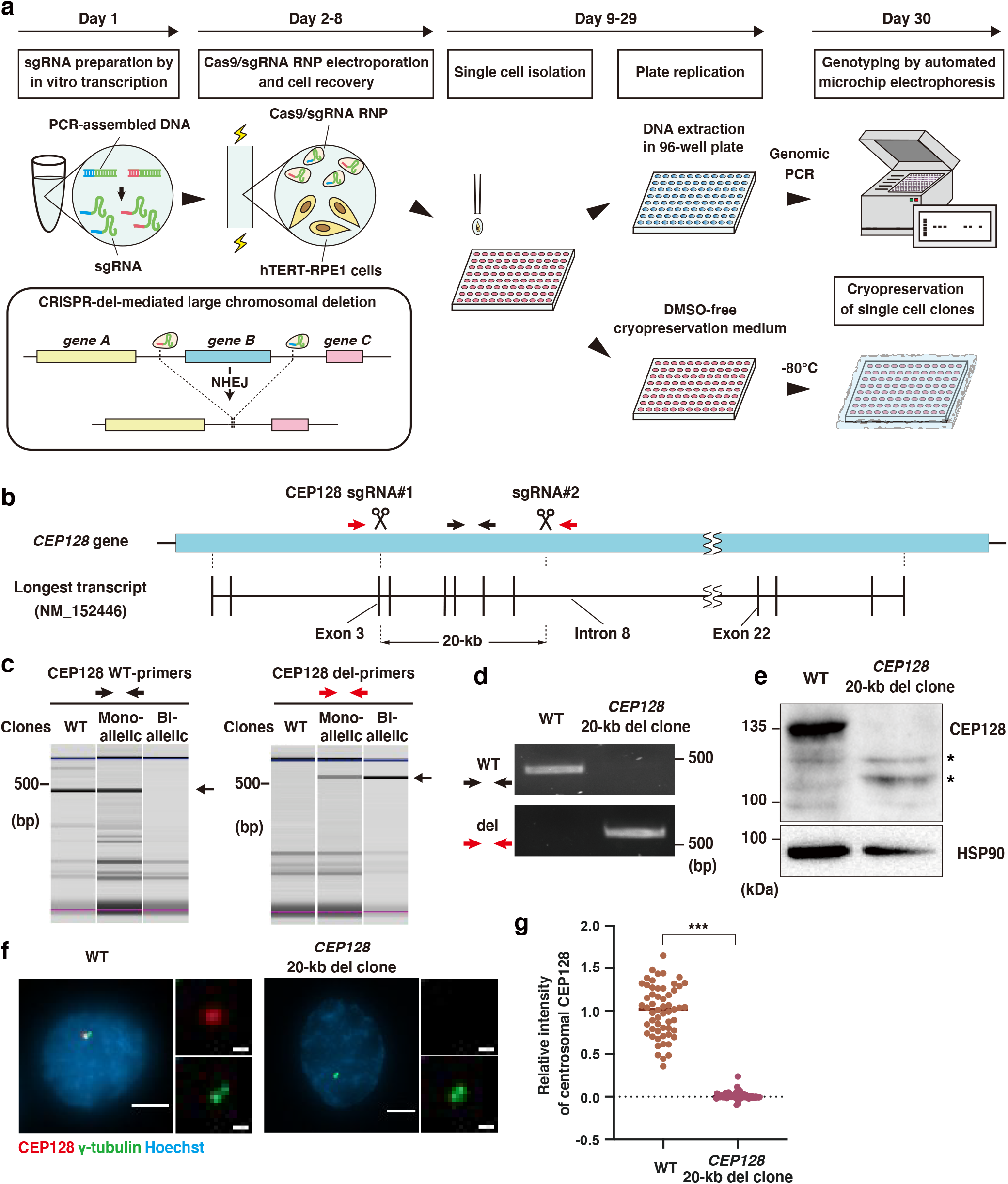
An optimized pipeline of CRISPR-del enables efficient gene knockout in human diploid RPE1 cells. **a**, Schematic overview of the optimized high-throughput CRISPR-del method. Cas9/sgRNA complexes introduce a large deletion of a chromosomal region flanked by the two sgRNA target sequences. **b**, Schematic representation of the *CEP128* gene and the longest transcript variant annotated in genome databases. The target positions of sgRNAs and the expected deletion length are shown. Black and red arrows indicate primers to detect WT and the deleted region of *CEP128* gene, respectively.**c**, Genomic PCR for detection of WT and the 20-kb deleted alleles of *CEP128* gene using the indicated primers, analyzed by the automated microchip electrophoresis system. Each electrophoresis pattern was adjusted according to the upper (blue) and lower (pink) size makers. The arrows on the side of electrophoresis images indicate the specific PCR product. **d**, Genomic PCR as in (**c**) with purified genomic DNA, analyzed by agarose gel electrophoresis. **e,**Western blotting to analyze the protein expression of CEP128 in WT cells and a *CEP128* 20-kb deleted clone. HSP90 was used as loading control. Asterisks show smaller fragments of CEP128 protein. **f**, Immunofluorescence imaging of CEP128 and γ-tubulin in WT cells and the 20-kb deleted clone. Scale bar: 5 μm (1 μm for insets). **g**, Quantification of relative CEP128 intensity at the centrosome from (**f**), ≥50 cells each. Data are represented as mean and *P* value was calculated by Mann–Whitney U test. ***P < 0.001.

To prove that our optimized CRISPR-del pipeline can be used for gene knockout in RPE1 cells, we applied it to target the *CEP128* gene, which encodes a centrosomal protein. The length of this gene is 502.54-kb, which is longer than 95% of the human protein-coding genes (Soheili-Nezhad, 2017). We first designed two sgRNAs (named as CEP128 sgRNA#1 and #2) to delete a 20-kb region ranging from 35 bp downstream of the first ATG in exon 3 to the middle of intron 8 of the longest CEP128 transcript variant (NM_152446) (Fig. 1b). Electroporation of Cas9 protein with the combination of sgRNA#1 and #2, but not with each individual, resulted in a successful chromosomal deletion (Fig. S1a). We then performed genotyping analysis of single cell clones from the *CEP128*-deleted cell pool. Microtip electrophoresis analyses followed by direct genomic PCRs revealed one clone, in which a deletion, but not a WT DNA band was detected, suggesting both alleles of the gene were edited (Fig. 1c). Further genotyping using the purified genomic DNA of this clone confirmed the result, indicating that it has the expected 20-kb deletion in both *CEP128* alleles (Fig. 1d). Western blotting with an antibody against the CEP128 C-terminus region revealed that the full-length protein is not expressed in the mutant clone (Fig. 1e). Unexpectedly, smaller fragments recognized by the CEP128 antibody were specifically expressed in the 20-kb del clone. Since the fragments disappeared upon CEP128 knockdown (Fig. S1b), we conclude that they are smaller CEP128 fragments emerged probably due to alternative translation initiation or nonsense-associated altered splicing on the deleted CEP128 transcripts in the mutant clone. Nevertheless, the absence of both CEP128 and its downstream protein Centriolin was confirmed at the centrosome of the clone by immunofluorescent analyses (Fig. 1f, 1g, S1c and S1d). Taken together, these data indicate that the optimized CRISPR-del pipeline can be used for successful gene knockout in human diploid RPE1 cells.

### The CRISPR-del pipeline effectively generates a nearly complete, bi-allelic deletion of a large protein-coding gene in RPE1 cells

For the purpose to achieve complete gene knockout in human cells, it would be important to estimate the length of the genomic DNA that can be eliminated by the current genome editing technology. Accordingly, we analyzed of what length and how frequently genomic deletions could be achieved in RPE1 cells with the CRISPR-del pipeline for routine lab work. Since the full length of *CEP128* is 502.54 kb, we tried to introduce deletions larger than the successfully obtained 20-kb one. In combination with sgRNA#1 that targets the site around the first ATG in exon 3, two other sgRNAs were designed at the downstream to delete 50-kb, 200-kb and 440-kb of the gene, respectively (Fig. 2a). For all deletions, the more effective one of the two sgRNAs was used in combination with sgRNA#1 to quantify their deletion frequencies (Fig. S2a and b). Single cell clones from two 96-well plates were genotyped for all three conditions (Fig. S2c). In the case of the targeted 50-kb deletion, we found 132 (69.8%) and 6 (3.2%) clones harboring mono- and bi-allelic deletions respectively among 189 surviving clones (Fig. 2b). For 200-kb and 440-kb targeted deletions, the frequency of the mono-allelic deletion was calculated as 42.0% (74/176) and 26.6% (45/169), respectively. Even though the frequency of bi-allelic deletions was also reduced as the deletion length increases (200-kb; 5/176, 2.8%), 3 out of 169 clones (1.8%) were found to have the very long 440-kb deletion for both alleles. Considering the number of deleted alleles out of the total number of alleles of all analyzed clones combined, deletion frequency for 50-kb, 200-kb and 440-kb were calculated to be 38.1%, 23.9%, 15.1%, respectively (Fig. 2b). The relationship between the length and the frequency of chromosomal deletion was an inverse correlation (Fig. 2c). Compared to the mono-allelic deletions, the frequency of the bi-allelic deletions was very low in this experiment. This may imply that the bi-allelic knockout of CEP128 reduces the proliferative ability of the cells. The genotype of the 440-kb deleted clones was confirmed by genomic PCR with the purified DNA (Fig. 2d) and subsequent genomic sequencing (Fig. 2e and Fig. S2d). Consistently, Western blotting showed that the expression of CEP128 was abolished in these clones (Fig. 2f). Immunofluorescence microscopy further confirmed that the downstream protein Centriolin was absent from the centrosome of the mutant clone (Fig. S1c and d). These analyses demonstrate that successful deletions were achieved for almost an entire locus in both *CEP128* alleles in RPE1 cells. In the human genome more than 95% of the protein-coding sequences have a length shorter than the here demonstrated possible deletion of 440 kb (Soheili-Nezhad, 2017). Our data therefore indicate that most of the human genes could be completely deleted by the CRISPR-del pipeline.

**Fig.2:**
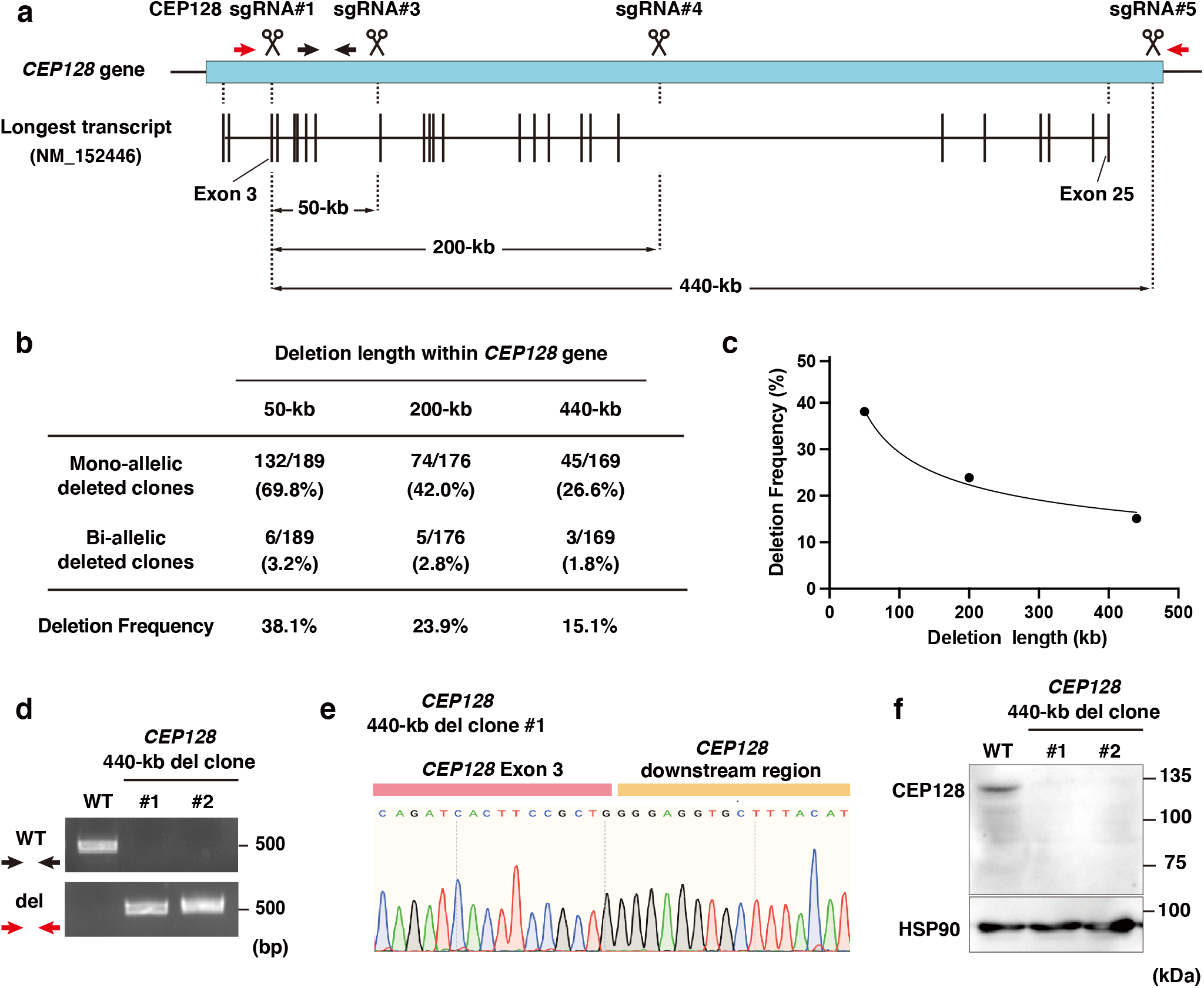
Analysis of efficacy in mono- and bi-allelic gene knockouts with a variety of deletion lengths using the CRISPR-del pipeline. **a**, Schematic representation of *CEP128* gene and the longest transcript variant annotated in genome databases. The target positions of sgRNAs and the expected lengths of large deletions are shown. Black and red arrows indicate primers to detect WT and the deleted regions, respectively. **b**, Summary for the efficiency of mono- and bi-allelic deletions within *CEP128* gene. **c**, A graph showing a relationship between the length and the frequency of chromosomal deletion from **c**. **d**, Genomic PCR for detection of WT and the 440-kb deleted alleles of *CEP128* gene using the indicated primers. **e**, Sequencing result of the CEP128 deleted alleles in the 440-kb deleted clone #1. **f**, Western blotting to analyze the protein expression of CEP128 in the lysate of WT cells and the 440-kb deleted clones. HSP90 was used as loading control.

### A high-throughput quantification reveals highly efficient large chromosomal deletions by the optimized CRISPR-del

To further analyze the deletion efficiency in a high-throughput and quantitative manner, we engineered a knock-in cell pool expressing a fluorescent protein and designed a FACS-based experimental system that can determine whether or not gene deletion has occurred based on the presence or absence of fluorescence. Using the CRISPR/Cpf1 system and a PCR-assembled dsDNA donor template, the nuclear ribonucleoprotein-coding *HNRNPA1* gene was targeted for endogenous tagging with mNeonGreen (mNG) at its C-terminus via homologous recombination (Fig. 3a). After genome editing, cells having mNG signal were enriched by cell sorting (Fig. S3a). Fluorescence imaging analysis revealed a specific nuclear signal of mNG in most of the genome-engineered cells (Fig. 3b and S3a). By genomic PCR, the proportions of cells with mono- and bi-allelic knock-ins in the cell pool were estimated to be approximately 62% and 38%, respectively (Fig. S3b-d). To verify the quantification strategy, we first tried to remove a 20-kb region containing the mNG sequence inserted to the *HNRNPA1* gene. For this purpose, we designed the first sgRNA (named as sgRNA#0) to target a sequence in *NFE2*, which is an upstream gene of *HNRNPA1*,and two versions of the second sgRNAs (named as sgRNA 20-kb#1 and #2) to target the intron 7 or 8 of *HNRNPA1* respectively (Fig. 3a). We then performed CRISPR-del in RPE1 *HNRNPA1-mNG* cells using these different combinations of sgRNAs. Ten days after the electroporation, we found that the combination of sgRNA#0 with sgRNA 20-kb#2, but not with a non-specific control sgRNA (CTRL), resulted in the appearance of a certain number of cells with significantly reduced mNG signal in the nucleus (Fig. 3b). This data shows that the reduction of the mNG signal can be used to monitor CRISPR-del efficiency.

**Fig.3:**
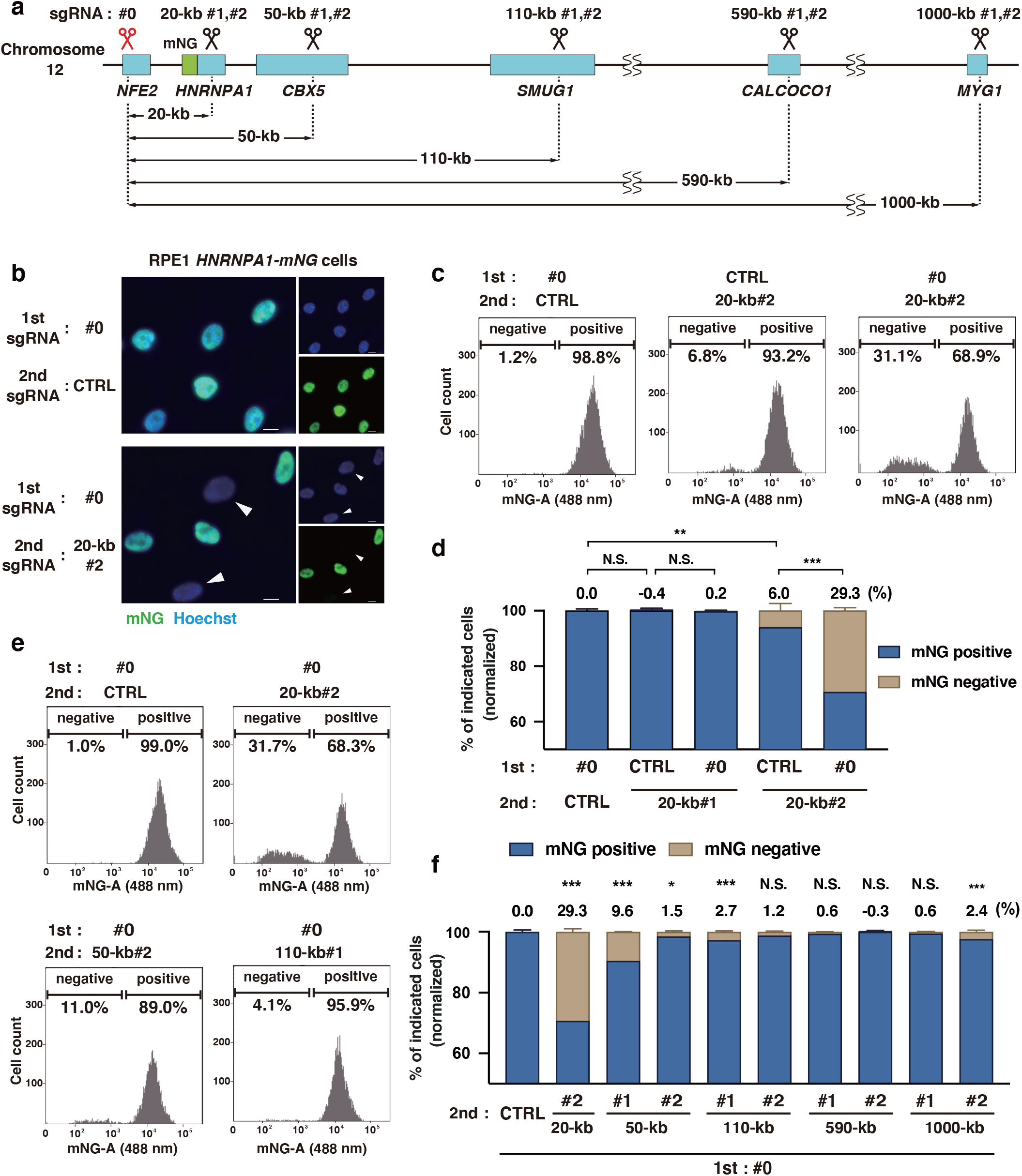
Quantitative analyses of length-dependent DNA deletion efficiency for the optimized CRISPR-del method using flow cytometry. **a**, Schematic representation of the chromosome region around the *HNRNPA1* gene locus. The mNG tag was inserted into the chromosomal site at the C-terminus of *HNRNPA1* gene. The target positions of sgRNAs and the expected lengths of large deletions are shown. **b**, Fluorescence imaging of HNRNPA1-mNG in RPE1 *HNRNPA1-mNG* cells electroporated with Cas9 protein and the indicated sgRNA pairs. Scale bar: 10 μm. **c,**FACS analyses for the mNG expression in RPE1 *HNRNPA1-mNG* cells electroporated with the indicated sgRNAs as in (b). Cells at 10 days after Cas9/sgRNAs electroporation were analyzed. Arrowheads indicate cells without the expression of HNRNPA1-mNG. **d**, Quantification of mNG positive and negative cells for each sgRNA pair from (c). The mean percentage of mNG-negative cells in control (1.47 %) was subtracted from each data. N = three biologically independent samples. 5000 cells were analyzed for each sample. The percentage of mNG-negative cells for each sample is indicated on top of the histogram. **e**, FACS analyses with the indicated conditions, as in (c). **f**, Quantification of (e), as in (d). Data are represented as mean ± S.D. and *P* value was calculated by Tukey–Kramer test. *P < 0.05, **P < 0.01, ***P < 0.001, N.S.: Not significant.

For high-throughput quantification of the gene deletion efficiency, the RPE1 *HNRNPA1-mNG* cells subjected to CRISPR-del were analyzed for fluorescent signal by a high-content flow cytometer. For the control sgRNA pair (sgRNA#0 and sgRNA CTRL), 1.5 ± 0.6% of the cells were not expressing the mNG protein (Fig. 3c and d). Since the cell line was not derived from a single clone, in this very small fraction of the cells, mNG might not have been properly knocked in. This percentage was therefore subtracted from that of the mNG negative cells in the other samples. In the case of sgRNA 20-kb#1, the combination with sgRNA#0 had no effect on the population of mNG-negative cells. For the experiments in this figure, the designed sgRNAs were directly used without any validations of genome editing efficiency except for sgRNA#0. Thus, sgRNA 20-kb#1 is rendered a non-effective sgRNA. Unexpectedly, another control combination (sgRNA CTRL and sgRNA 20-kb#2) resulted in the production of mNG negative cells at 6 ± 2.6%, although the sgRNA 20-kb#2 targets an intron sequence of *HNRNPA1*. This targeted intron region might be critical for the HNRNPA1 expression. Nevertheless, the combination for 20-kb deletion (sgRNA#0 and sgRNA 20-kb#2) significantly increased the ratio of mNG-negative cells to 29.3 ± 1.1%. Given that HNRNPA1 is an abundant protein due to its strong housekeeping promoter (Biamonti et al., 1993), most of the cells without mNG signal likely have the deletion of the target region. Taken together, these data indicate that a 20-kb chromosomal deletion could be achieved with more than 20% frequency in RPE1 cells by using the optimized CRISPR-del.

We further analyzed the gene deletion efficiency for much longer genomic regions by the optimized CRISPR-del. To avoid closed chromatin regions, second sgRNAs were designed to target sequences within downstream genes of *HNRNPA1*.The second sgRNAs for the deletion of 50-kb, 110-kb, 590-kb and 1000-kb targeted *CBX5, SMUG1, CALCOCO1* and *MYG1* genes, respectively. In combination with the first sgRNA#0, the effect of each second sgRNA for gene deletion efficiency was quantitatively analyzed as mentioned above. Despite the variable results among the sgRNA combinations, at least one of the two combinations significantly increased the ratio of mNG-negative cells, compared to the control, for all conditions except for 590-kb (Fig. 3e and f). Surprisingly, an increase of mNG-negative cells to 2.4 ± 0.6% was observed with a sgRNA combination targeting a 1000-kb deletion. By genomic PCR (Fig. S3e and f) and subsequent genomic sequencing (Fig. S3g), the occurrence of the 1000-kb deletion was confirmed in the purified DNA from the cells electroporated with Cas9 and the sgRNA pair. Collectively, these quantitative analyses revealed that, using effective sgRNA pairs, our optimized CRISPR-del enables deletion of a large chromosome region in RPE1 cells at high frequency.

### The optimized CRISPR-del can introduce a homozygous deletion of cancer-associated large genomic regions into human diploid cells

Large chromosomal deletions are frequently observed in human cancers (Beroukhim et al., 2010; Bignell et al., 2010). To investigate the impact of chromosome deletions on the development and progression of cancers, it is important to create model cell lines bearing these cancer-associated hetero- or homozygous deletions. The 9p21.3 region, which contains the three tumor suppressor genes *CDKN2A, CDKN2B*, and *MTAP*, is known to be frequently deleted in different types of cancer cells (Baker et al., 2016; Frigerio et al., 2014; Sasaki et al., 2003). Especially in bladder cancer cell lines, this region is commonly lost and the deletions are extended to the up- and downstream genes of the tumor suppressors (Jinek et al., 2012). Therefore, we tested whether the optimized CRISPR-del can introduce a homozygous deletion, which is similar to that identified in the bladder cancer cell line UM-UC-3 (Jinek et al., 2012), into the genome of non-transformed RPE1 cells (Fig. 4a). To delete the 650-kb region in chromosome 9, first and second sgRNAs were designed to target an upstream sequence of *MTAP* and a downstream sequence of *DMRTA1*. After CRISPR-del editing followed by single cell cloning, genotyping was performed using different primer pairs to detect the four individual genes in the region. We identified a clone lacking all four genes (Fig. 4b) and showing a specific PCR product indicating the expected large deletion (Fig. 4c). Western blotting confirmed that p15, a protein encoded by *CDKN2B*, was not expressed in the 9p21.3 large deletion clone (Fig. 4d). These results demonstrate that our optimized CRISPR-del pipeline can be applied for the generation of model cell lines with cancer-associated large chromosomal deletions in a non-transformed human diploid cell background.

**Fig.4:**
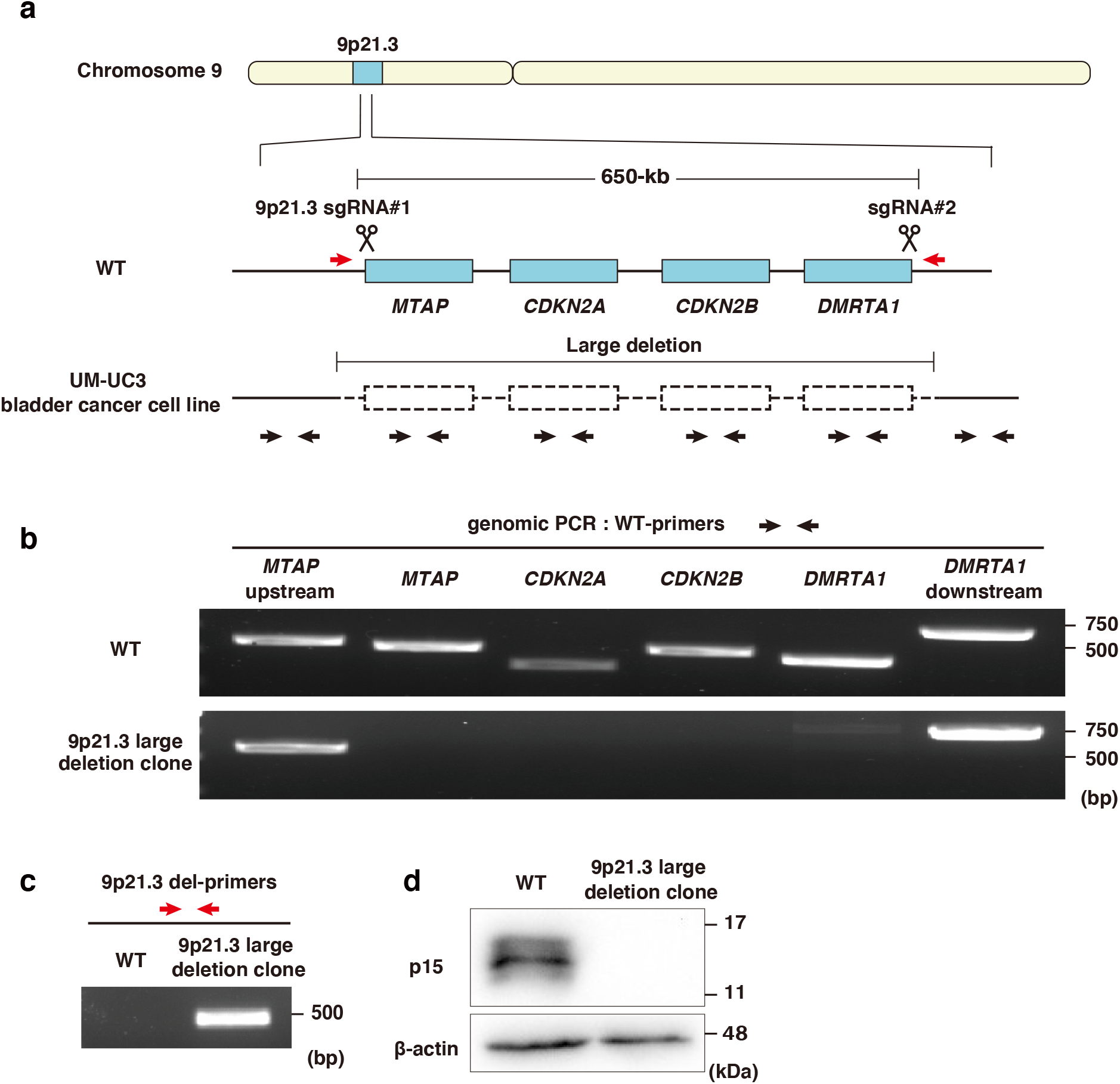
Mimicking a cancer-associated large chromosome deletion by using the CRISPR-del pipeline in non-transformed human cells. **a**, Schematic representation of the chromosome 9p21.3 region in the human genome. UM-UC-3 bladder cancer cell line intrinsically has a large deletion containing the *MTAP, CDKN2A, CDKN2B, DMRTA1* genes. The target positions of sgRNAs are shown. Black and red arrows indicate primers to detect WT and deleted genomic regions, respectively. **b**, Genomic PCR for detection of the indicated regions within 9p21.3 in WT cells and a 9p21.3 large deletion clone. **c**, Genomic PCR for detection of the large deletion in WT cells and the 9p21.3 large deletion clone. **d**, Western blotting to analyze the protein expression of p15, the *CDKN2B* gene product, in WT cells and the deletion clone. β-actin was used as loading control.

## Discussion

The indel-based gene editing via the CRISPR/Cas9 technology is a well-established method for a simple, easy and cost-effective gene disruption, as compared to previous genome editing techniques. Despite its wide usage and benefits, this method is not so reliable for “complete” gene knockout, as several mechanisms can function to nullify the effects of indels. In this study, we established a pipeline of CRISPR-del, which can overcome the drawbacks of the indel-based method by removing an entire gene locus, for effective use in a non-transformed human diploid RPE1 cell line.

Although the indel-based gene disruption is considered a generally simple method, the genotyping procedure for selecting knockout clones is time consuming and labor intensive due to the necessity for detection of these short indels. On top of that, the standard assays for the indel detection, such as the sequencing of genomic PCR products and the Surveyor assay using a mismatch specific nuclease, are costly. In contrast, the genotyping for CRISPR-del consists only of the preparation of genomic PCRs for detection of the WT and deleted alleles. In addition to this simple and cost-effective genotyping, our optimized pipeline of CRISPR-del includes several other fine-tuned steps: 1) the synthesis of sgRNAs from PCR-assembled DNA templates by *in vitro* transcription enables a cloning-free genome editing; 2) the electroporation of Cas9/sgRNA RNPs gives high editing efficiency with low off-target effects; 3) the use of an automated single cell dispensing system based on piezo-acoustic technology provides an efficient single cell cloning with high viability; 4) DMSO-free cryopreservation medium allows for a direct and mild freezing of cells in 96-well plates without removing cell dissociation solution; 5) the genomic DNA is directly extracted from single cell clones in 96-well plates for genotyping PCR, and 6) the PCR products are automatically analyzed by a microtip electrophoresis system. These optimizations improve the CRISPR-del approach in terms of efficacy and cost, and allow for it to be implemented in high-throughput gene knockout studies of human diploid cells.

CRISPR-del has been considered to be a technique with a relatively low probability of successful knockout, as it was previously reported that 65% of sgRNA combinations yield deletion efficiency of <20% (Canver et al., 2014). However, in this study we implement optimizations to the method that demonstrates a much higher deletion rate. Quantitative analyses revealed that the optimized method successfully deleted chromosomal regions of 50-kb, 200-kb, and 440-kb with 38.1%, 23.9%, and 15.1% efficiency, respectively. Although a fair comparison of the deletion efficiency between our approach and that of the previously described CRISPR-del methods is difficult due to the different quantitative analyses performed (He et al., 2015; Li et al., 2021), the study by Canver et al. calculated this factor in a similar manner (Canver et al., 2014). The authors conducted CRISPR-del in murine erythroleukemia cells, which led to chromosomal deletions of 20-kb, 71-kb and 1026-kb, with deletion efficiencies of 24.3%, 0.3% and 0.7%, respectively (Canver et al., 2014). Despite the poor transfectability of RPE1 cells, we demonstrate that our optimizations have dramatically improved CRISPR-del, rendering it a more effective method for large chromosomal deletions.

Based on the past experiments using CRISPR-del, the correlation between size and frequency of targeted chromosomal deletions seems to be controversial (Canver et al., 2014; He et al., 2015; Li et al., 2021). Our quantitative analyses show an inverse relationship between these parameters, similar to the previous report (Canver et al., 2014). Although the length of the chromosomal deletion that can be introduced is a critical factor in performing a “complete” gene knockout by CRISPR-del, our optimized method successfully achieved deletions of very long regions of genomic DNA: 440-kb for the *CEP128* gene, 1000-kb for the region around the *HNRNPA1* gene and 650-kb for the chromosomal region 9p21.3. This range covers the length of more than 95 % of the human genes encoding proteins (Soheili-Nezhad, 2017), indicating that the CRISPR-del pipeline can be used to generate “complete” knockout cell lines for most of the human protein-coding genes as routine lab work.

## Acknowledgements

We thank the Kitagawa lab members and Dr. Elmar Schiebel from ZMBH in Heidelberg University for technical supports and helpful discussions. This work was supported by JSPS KAKENHI grants (Grant numbers: 18K06246, 19H05651, 20K15987, 20K22701, 21H02623) from the Ministry of Education, Science, Sports and Culture of Japan, the PRESTO program (JPMJPR21EC) of the Japan Science and Technology agency, Takeda Science Foundation, Mochida Memorial foundation, Daiichi Sankyo Foundation of Life Science, The Uehara Memorial Foundation and The Research Foundation for Pharmaceutical Sciences.

## Author contributions

S.H. conceived and designed the study. T.K. performed most of the experiments. A.M. generated the knock-in cell pool. M.G. optimized the genome editing conditions. T.H. performed clone validation, M.F. and T.C. provided suggestions. T.K., S.H. and D.K. analyzed the data. S.H. and D.K. wrote the manuscript with input from T.K. All authors contributed to discussions and manuscript preparation.

## Material and methods

### Cell culture

RPE1 cells were grown in Dulbecco’s Modified Eagle’s Medium F-12 (DMEM/F-12) with 10% FBS and 1% penicillin/streptomycin. All cell lines were cultured at 37°C in a humidified 5% CO2 incubator.

### sgRNA synthesis

sgRNAs were transcribed *in vitro* from PCR-generated DNA templates according to a previously published method (DeWitt et al., 2016) with slight modifications. Briefly, template DNA was assembled by PCR from five different primers: 1) a variable forward primer containing T7 promoter and desired guide sequence, 2) a variable reverse primer containing the reverse complement of the guide sequence and the first 15 nt of the non-variable region of the sgRNA, 3) a forward primer containing the entire invariant region of the sgRNA, and 4), 5) two amplification primers. The assembled template was purified and subjected to *in vitro* transcription by T7 RNA polymerase using the Hiscribe T7 High Yield RNA Synthesis Kit (New England Biolabs). The reaction product was treated with DNase I, and the synthesized sgRNA was purified using the RNA Clean & Concentrator Kit (ZYMO RESEARCH). All sgRNA and primer sequences used in this study are listed in Supplementary Table1 and 2, respectively.

### CRISPR-del-mediated gene knockout

For large chromosomal deletions, the CRISPR-del method was performed with Cas9 protein and two synthesized sgRNAs. Cas9/sgRNA RNPs were electroporated into RPE1 cells stably expressing Tet3G transactivator (described as “RPE1 cells” in this study) using the Neon Transfection System (Thermo Fischer Scientific) according to the manufacturer’s protocol. Briefly, HiFi Cas9 protein (1.55 μM) from Integrated DNA Technologies (IDT) and two sgRNAs (0.92 μM each) were pre-incubated in Resuspension buffer R and mixed with cells (0.125 x10^5^/μl) and Cas9 electroporation enhancer (1.8 μM, IDT). After resuspension, electroporation was immediately conducted using a 10 μl Neon tip at a voltage of 1300 V with two 20 ms pulses. The transfected cells were seeded into a 24-well plate. After recovery from the electroporation, single cells were isolated into 96-well plates using cellenONE (cellenion) according to the manufacturer’s protocol. After cell expansion, each 96-well plate was duplicated for genotyping and preparation of a frozen stock. Briefly, cells in 96-well plates were washed with PBS and treated with 25 μl of Trypsin/EDTA solution (nacalai tesque). After brief incubation at 37°C, the detached cells were resuspended with 75 μl of Cell Reservoir One (nacalai tesque), a DMSO-free cryopreservation medium. 25 μl of the cell mixture were transferred into a well of another 96-well plate filled with 175 μl of growth medium for cell expansion followed by genotyping analysis. The 96-well plate with the remaining cell suspension was placed in a deep freezer at - 80°C.

### Genotyping

For high-throughput genotyping, genomic DNA was directly extracted from single cell clones in 96-well plates using DNAzol Direct (Molecular Research Center) and subjected to PCR for the detection of both WT and the deleted alleles using appropriate primers. Briefly, after removal of culture medium, 20 μl of DNAzol Direct was added to each well and the 96-well plate was shaken at 800 rpm for 20 min at room temperature. 1μl of the lysate containing genomic DNA was used for 10 μl of PCR reaction using KOD One PCR Master Mix (TOYOBO). The PCR products were analyzed by the automated microchip electrophoresis system MCE-202 MultiNa (Shimadzu). To confirm the genotype of homozygous KO clones, their genomic DNA was purified using NucleoSpin DNA RapidLyse kit (Macherey-Nagel) and subjected again to genotyping PCR. The PCR products were analyzed by agarose gel electrophoresis and sanger sequencing.

### CRISPR/Cpf1-mediated gene knock-in

Endogenous mNG tagging of HNRNPA1 by CRISPR-Cpf1 system was performed with the electroporation of Cpf1/crRNA RNP and dsDNA repair template. crRNA was designed to target the site immediately downstream of the stop codon of *HNRNPA1* and transcribed *in vitro* as described above. The DNA template for crRNA synthesis was assembled by PCR using a forward primer containing T7 promoter and the target sequence, and a reverse primer containing the reverse complement of the target sequence and the non-variable region of crRNA. The dsDNA repair template was amplified by PCR from a plasmid encoding the mNG sequence using two primers containing 90 bp left and right homology arm sequence, respectively. Electroporation of Cpf1/crRNA and the repair template was conducted similarly to the Cas9/sgRNA condition described above, with a modification in the electroporation solution. A.s.Cpf1 Ultra (1 μM, IDT) and crRNA (1 μM) were pre-incubated in buffer R and mixed with RPE1 cells (0.125 ×10^5^ /μL), Cpf1 electroporation enhancer (1.8 μM, IDT) and the repair template (33 nM). mNG-positive cells were sorted using FACS Aria III (BD Biosciences), equipped with 355/405/488/561/633 nm lasers. After cell expansion, cell sorting was repeated to re-enrich mNG-positive cells. The knock-in cell pool was subjected to CRISPR-del experiments. The crRNA sequence is listed in Supplementary Table1.

### CRISPR-del efficiency assessed by Flow cytometry

HNRNPA1-mNG knock-in cells were electroporated with Cas9 and the indicated sgRNA pairs for CRISPR-del application. After 10 days in culture, the cells were harvested with trypsin/EDTA solution, washed in cold PBS, and analyzed by FACS Aria III for mNG expression. Data for approximately 5,000 gated events were collected. Due to the presence of a very small portion of mNG-negative cells in control samples, the mean percentage (1.47 %) was then subtracted from that of each CRISPR-del sample before comparison.

### siRNA-mediated gene knockdown

Lipofectamine RNAiMAX (Thermo Fisher) was used with a final concentration of 20 nM siRNA for siRNA transfection according to manufacturer’s protocol. Transfected cells were harvested at 72 h after transfection for Western blotting. Control siRNA (4390843) and CEP128 siRNA pool (L-032761-02) were purchased from Life Technologies and Dharmacon, respectively.

### Western blotting

Cells were lysed on ice in lysis buffer (20 mM Tris-HCl, pH 7.5, 50 mM NaCl, 1% Triton X-100, 5 mM EGTA, 1 mM DTT, 2 mM MgCl2, and 1:1,000 protease inhibitor cocktail [Nakarai Tesque]). After centrifugation, the supernatant was added to Laemmli sample buffer, boiled and subjected to SDS-PAGE. Separated proteins were transferred onto Immobilon-P membrane (Merck) using Trans-Blot SD Semi-Dry Electrophoretic Transfer Cell (Bio-Rad Laboratories). The membranes were probed with the primary antibodies, followed by incubation with their respective HRP-conjugated secondary antibodies (Promega). The membrane was soaked with Chemi-Lumi Super (Nakarai Tesque) for the signal detection using ChemiDoc XRS+ (Bio-Rad Laboratories).

### Immunofluorescence

For the immunofluorescence analyses, cells cultured on coverslips (Matsunami) were fixed with −20 °C methanol for 7 min, or 4 % PFA at room temperature for 30min. Fixed cells were incubated with blocking buffer (1 % bovine serum albumin in PBS containing 0.05 % Triton X-100) for 30 min at room temperature. The cells were then incubated with primary antibodies in the blocking buffer for 1 hour in a humid chamber. After washing with PBS, the cells were incubated with secondary antibodies and Hoechst 33258 (DOJINDO, 1:3000–1:5000) in the blocking buffer for 30 min, followed by final wash with PBS. The coverslips were mounted onto glass slides (Matsunami) using ProLong Gold (Molecular Probes), with the cell side down.

### Antibodies

The following primary antibodies were used in this study: anti-CEP128 (abcam, ab118797; IF 1:500, WB 1:1000), anti-γ-tubulin (Merck, GTU88; IF 1:1,000), anti-p15 (Santa Cruz Biotechnology, sc-271791; WB 1:1000), anti-HSP90 (BD Biosciences, 610419; WB 1:1000) and anti-*β*-actin (Santa Cruz Biotechnology, sc-47778; WB 1:1000). The following secondary antibodies were used: anti-mouse IgG Alexa Fluor 488 (Molecular Probes, 1:1000), anti-rabbit IgG Alexa Fluor 555 (Molecular Probes, 1:1000), anti-mouse IgG HRP (Promega, WB 1:10000) and anti-rabbit IgG HRP (Promega, WB 1:10000).

### Statistics analysis

Statistical comparison between the data from different groups was performed in PRISM v.8 software (GraphPad) using either a Mann–Whitney U test or a Tukey–Kramer test as indicated in the legend. P values <0.05 were considered statistically significant. All data shown are mean ± S.D. The sample size is indicated in the figure legends.

## Supplementary figure legends

**Fig.S1:**
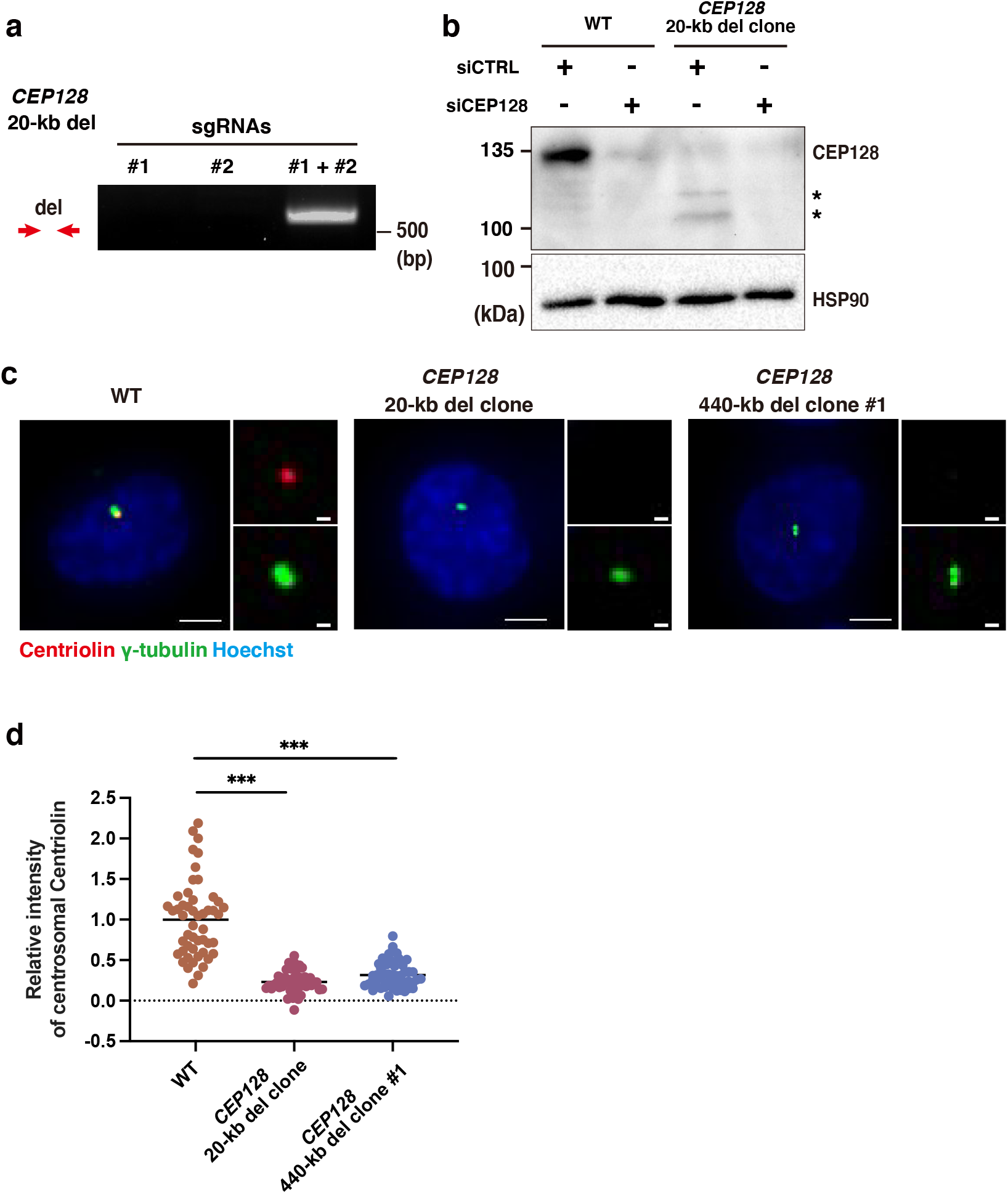
Validation of CEP128 deletion mutants. **a**, Genomic PCR to detect the 20-kb deletion in cells electroporated with Cas9 and the indicated sgRNAs. **b**, Western blotting to analyze the protein expression of CEP128 in the cell lysate of WT and a *CEP128* 20-kb deleted clone at 72 hr after transfection of the indicated siRNA. Asterisks show smaller fragments of CEP128 protein. **c**, Immunofluorescence imaging of Centriolin and γ-tubulin in the cells of WT, the 20-kb or 440-kb *CEP128*-deleted clone. Scale bar: 5 μm (1 μm for insets). **d**, Quantification of relative Centriolin intensity at the centrosome from (**a**), 50 cells for each sample. Data are represented as mean and *P* value was calculated by Mann–Whitney U test. ***P < 0.001.

**Fig.S2:**
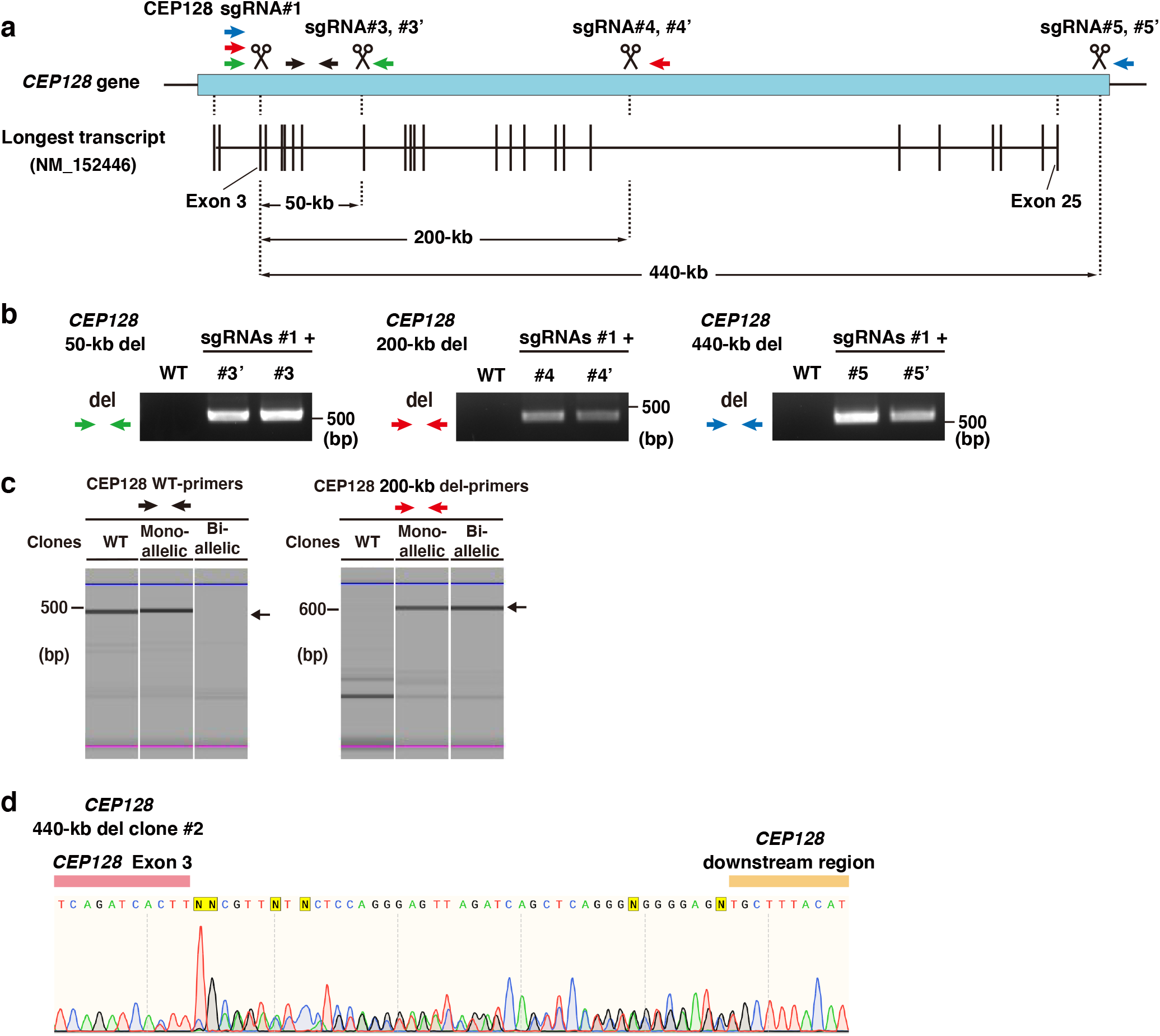
Large chromosomal deletions within the *CEP128* gene by the CRISPR-del pipeline. **a**, Schematic representation of *CEP128* gene and the longest transcript variant annotated in genome databases. The target positions of sgRNAs and the expected lengths of large deletions are shown. Green, red, blue and black arrows indicate primers to detect the 50-kb, 200-kb and 440-kb deleted and WT regions, respectively.**b**, For the validation of sgRNA combinations, chromosomal deletions with different lengths within the *CEP128* gene were detected by genomic PCR. CEP128 sgRNA#3, #4 and #5, together with sgRNA#1, were used for further analyses in Fig. 2 and Fig. S2. **c**, Genomic PCR for detection of WT and the 200-kb deleted alleles of *CEP128* gene, analyzed by the automated microchip electrophoresis system. Each electrophoresis pattern was adjusted according to the upper (blue) and lower (pink) size makers. The arrows on the right side of electrophoresis images indicate the specific PCR product. **d**, Sequencing result of the *CEP128* deleted alleles in the 440-kb deleted clone #2.

**Fig.S3:**
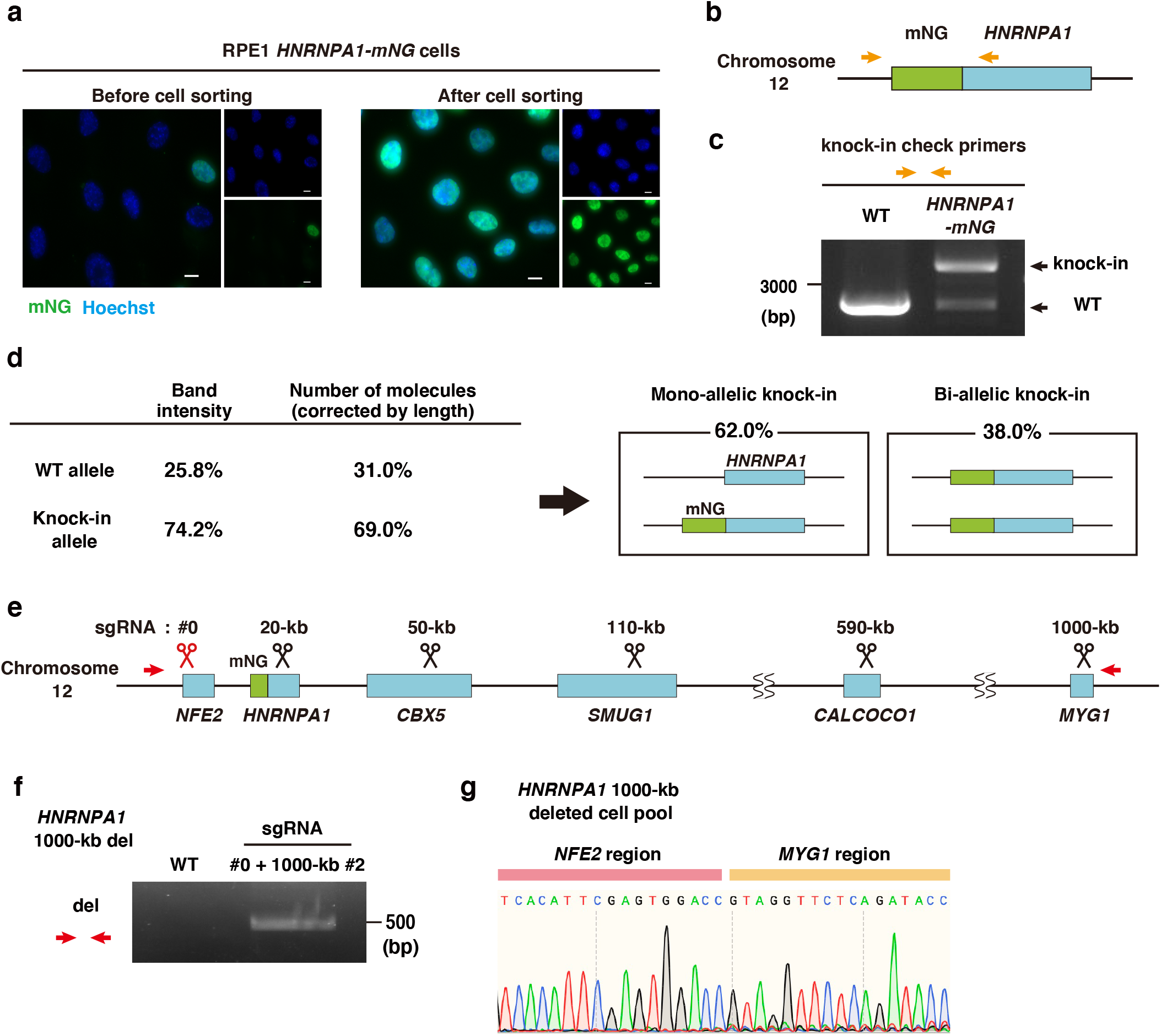
Large deletion of a chromosome region including *HNRNPA1* gene locus by the CRISPR-del. **a**, Fluorescence imaging of HNRNPA1-mNG in the pool of RPE1 *HNRNPA1-mNG* cells before and after enrichment of mNG-positive cells by cell sorting. **b**, Schematic representation of the *HNRNPA1* gene locus in RPE1 *HNRNPA1-mNG* cells. Orange arrows indicate the primers to detect the knock-in allele. **c**, Genomic PCR to detect WT and the knock-in alleles of the *HNRNPA1* gene in the cell pool. **d**, Summary for the proportions of cells with mono- and bi-allelic knock-ins in the cell pool from **c**. **e**, Schematic representation of the chromosome region around the *HNRNPA1* gene locus in RPE1 *HNRNPA1-mNG* cells. The target positions of sgRNAs are shown. Red arrows indicate the primers to detect the 1000-kb chromosomal deletion. **f**, Genomic PCR to detect the 1000-kb deletion in WT cells and cells electroporated with Cas9 protein and the indicated sgRNA pair. **g**, Sequencing result of the deletion band shown in (**c**).

