## Supplementary Table for "A CRISPR-del-based pipeline for complete gene knockout in human diploid cells"

**Komori T. *et al.* Supplementary Table 1**

| sgRNA/crRNA Target | Name | Sequence |
|---|---|---|
| CEP128 Exon3 | CEP128 sgRNA#1 | GACTCAATCGGTCACGACAG |
| CEP128 Intron8(20-kb) | CEP128 sgRNA#2 | GATTTTTTCGGATGTACCAA |
| CEP128 Intron8(50-kb) | CEP128 sgRNA#3 | CCTGACAAGGAACCTTATTG |
| CEP128 Intron19(200kb) | CEP128 sgRNA#4 | TCAAGAAGGGCACTTTACGT |
| CEP128 downstream region(440-kb) | CEP128 sgRNA#5 | GATGTATAGATAACACGGGG |
| NFE2 | HNRNPA1#0 | TCACATTCGAGTGGACCATC |
| HNRNPA1 | HNRNPA1 20-kb#1 | TGTGCAAGTGTAACGGCTGA |
| HNRNPA1 | HNRNPA1 20-kb#2 | GATTGGAGTCACCATAGTTT |
| CBX5 | HNRNPA1 50-kb#1 | ACCCAGGGAGCACAATACTT |
| CBX5 | HNRNPA1 50-kb#2 | TACCCAGGGAGCACAATACT |
| SMUG1 | HNRNPA1 110-kb#1 | GAAGTCTCTTATACCCACGG |
| SMUG1 | HNRNPA1 110-kb#2 | TGGGAACCATCCAATCCCT |
| CALCOCO1 | HNRNPA1 590-kb#1 | AAGGACCTGCACGCTCACCA |
| CALCOCO1 | HNRNPA1 590-kb#2 | CACTTGAGACAACCCACTCA |
| MYG1 | HNRNPA1 1000-kb#1 | TGGCACCTTCCACTGCGACG |
| MYG1 | HNRNPA1 1000-kb#2 | GGTATCTGAGAACCTACCTC |
| MTAP | 9p21.3 sgRNA#1 | AAGTAAGCAGTTCTCCCACG |
| DMRTA1 | 9p21.3 sgRNA#2 | TAGTGGATGTGGAGCCCAAA |
| HNRNPA1 | crRNA HNRNPA1 | ATTAGGTAAGTAAGCACCTT |

**Komori T. *et al.* Supplementary Table 2**

| Primer for PCR-assembled DNA templates | Name | Sequence |
|---|---|---|
| sgRNA amplification primers | Universal Design Primer_Fw | TTCTAATACGACTCACTATAG |
| | Universal Design Primer_Rv | AAAAGCACCGACTCGGTG |
| crRNA_tracrRNA | crRNA_tracrRNA | GTTTTAGAGCTAGAAATAGCAAGTTAAAATAAGGCTAGTCCGTTATCAACTTGAAAAAGTGGCACCGAGTCGGTGCTTTT |
| CEP128 sgRNA#1 | sgRNA_CEP128_#1_Fw | TTCTAATACGACTCACTATAGACTCAATCGGTCACGACAG |
| | sgRNA_CEP128_#1_Rv | TTCTAGCTCTAAAACCTGTCGTGACCGATTGAGTC |
| CEP128 sgRNA#2 | sgRNA_CEP128_#2_Fw | TTCTAATACGACTCACTATAGATTTTTTCGGATGTACCAA |
| | sgRNA_CEP128_#2_Rv | TTCTAGCTCTAAAACTTGGTACATCCGAAAAAATC |
| CEP128 sgRNA#3 | sgRNA_CEP128_#3_Fw | TTCTAATACGACTCACTATAGCCTGACAAGGAACCTTATTG |
| | sgRNA_CEP128_#3_Rv | TTCTAGCTCTAAAACCAATAAGGTTCCTTGTCAGG |
| CEP128 sgRNA#4 | sgRNA_CEP128_#4_Fw | TTCTAATACGACTCACTATAGTCAAGAAGGGCACTTTACGT |
| | sgRNA_CEP128_#4_Rv | TTCTAGCTCTAAAACACGTAAAGTGCCCTTCTTGA |
| CEP128 sgRNA#5 | sgRNA_CEP128_#5_Fw | TTCTAATACGACTCACTATAGATGTATAGATAACACGGGG |
| | sgRNA_CEP128_#5_Rv | TTCTAGCTCTAAAACCCCCGTGTTATCTATACATC |
| HNRNPA1#0 | sgRNA_#0_Fw | TTCTAATACGACTCACTATAGTCACATTCGAGTGGACCATC |
| | sgRNA_#0_Rv | TTCTAGCTCTAAAACGATGGTCCACTCGAATGTGAC |
| HNRNPA1 20-kb#1 | sgRNA_del20-kb_#1_Fw | TTCTAATACGACTCACTATAGTGTGCAAGTGTAACGGCTGA |
| | sgRNA_del20-kb_#1_Rv | TTCTAGCTCTAAAACTCAGCCGTTACACTTGCACAC |
| HNRNPA1 20-kb#2 | sgRNA_del20-kb_#2_Fw | TTCTAATACGACTCACTATAGATTGGAGTCACCATAGTTT |
| | sgRNA_del20-kb_#2_Rv | TTCTAGCTCTAAAACAAACTATGGTGACTCCAATC |
| HNRNPA1 50-kb#1 | sgRNA_del50-kb_#1_Fw | TTCTAATACGACTCACTATAGACCCAGGGAGCACAATACTT |
| | sgRNA_del50-kb_#1_Rv | TTCTAGCTCTAAAACAAGTATTGTGCTCCCTGGGTC |
| HNRNPA1 50-kb#2 | sgRNA_del50-kb_#2_Fw | TTCTAATACGACTCACTATAGTACCCAGGGAGCACAATACT |
| | sgRNA_del50-kb_#2_Rv | TTCTAGCTCTAAAACAGTATTGTGCTCCCTGGGTAC |
| HNRNPA1 110-kb#1 | sgRNA_del110-kb_#1_Fw | TTCTAATACGACTCACTATAGAAGTCTCTTATACCCACGG |
| | sgRNA_del110-kb_#1_Rv | TTCTAGCTCTAAAACCCGTGGGTATAAGAGACTTC |
| HNRNPA1 110-kb#2 | sgRNA_del110-kb#_2_Fw | TTCTAATACGACTCACTATAGTGGGAACCATCCAATCCCT |
| | sgRNA_del110-kb#_2_Rv | TTCTAGCTCTAAAACAGGGATTGGATGGTTCCCAC |
| HNRNPA1 590-kb#1 | sgRNA_del590-kb_#1_Fw | TTCTAATACGACTCACTATAGAAGGACCTGCACGCTCACCA |
| | sgRNA_del590-kb_#1_Rv | TTCTAGCTCTAAAACTGGTGAGCGTGCAGGTCCTTC |
| HNRNPA1 590-kb#2 | sgRNA_del590-kb_#2_Fw | TTCTAATACGACTCACTATAGCACTTGAGACAACCCACTCA |
| | sgRNA_del590-kb_#2_Rv | TTCTAGCTCTAAAACTGAGTGGGTTGTCTCAAGTGC |
| HNRNPA1 1000-kb#1 | sgRNA_del1000-kb_#1_Fw | TTCTAATACGACTCACTATAGTGGCACCTTCCACTGCGACG |
| | sgRNA_del1000-kb_#1_Rv | TTCTAGCTCTAAAACCGTCGCAGTGGAAGGTGCCAC |
| HNRNPA1 1000-kb#2 | sgRNA_del1000-kb_#2_Fw | TTCTAATACGACTCACTATAGGTATCTGAGAACCTACCTC |
| | sgRNA_del1000-kb_#2_Rv | TTCTAGCTCTAAAACGAGGTAGGTTCTCAGATACC |
| 9p21.3 sgRNA#1 | sgRNA_9p21.3_#1_Fw | TTCTAATACGACTCACTATAGAAGTAAGCAGTTCTCCCACG |
| | sgRNA_9p21.3_#1_Rv | TTCTAGCTCTAAAACCGTGGGAGAACTGCTTACTTC |
| 9p21.3 sgRNA#2 | sgRNA_9p21.3_#2_Fw | TTCTAATACGACTCACTATAGTAGTGGATGTGGAGCCCAAA |
| | sgRNA_9p21.3_#2_Rv | GTAGTGGATGTGGAGCCCAAAGTTTTAGAGCTAGAA |
| HNRNPA1 | HNRNPA1_crRNA_Fw | TTCTAATACGACTCACTATAGTAATTTCTACTCTTGTAGAT |
| | HNRNPA1_crRNA_Rv | AAGGTGCTTACTTACCTAATATCTACAAGAGTAGAAATTAC |

| dsDNA repair template | Name | Sequence |
|---|---|---|
| HNRNPA1 | HNRNPA1_dsDNA repair template_Fw | CACTTTGAAACTTTAAAAGAAAAATTGTACTTTTCAGGTGGCTATGGCGGTTCCAGCAGCAGCAGTAGCTATGGCAGTGGCAGAAGAT TTGGAGCTGGTGCAGGTGCAG |
| | HNRNPA1_dsDNA repair template_Rv | ACTGCAATTATAATGTTAACTATGTTGCACTGCTCAGCTACATTAGGGTTATTGGGTTCATCAGCAATTTAAAAAATTATGTCAACACAC AAAAAGGTGCTTACTTACCTAACTACTTGTACAGCTCGTCCATGC |

| Primer for deletion check | Name | Sequence |
|---|---|---|
| CEP128 del-primers_20-kb | del-primer_CEP128_20-kb_Fw | AATGTTCTTAATGGGACCGTCT |
| | del-primer_CEP128_20-kb_Rv | GGGAGAATTACATGGGGAAGTGA |
| CEP128 del-primers_50-kb | del-primer_CEP128_50-kb_Fw | TGGCAGAATCATCCAGCGAATC |
| | del-primer_CEP128_50-kb_Rv | ATTATCCATGAGCATGACCTGCC |
| CEP128 del-primers_200-kb | del-primer_CEP128_200-kb_Fw | TGGCAGAATCATCCAGCGAA |
| | del-primer_CEP128_200-kb_Rv | GGAGATCCAAGGCAAGCACT |
| CEP128 del-primers_440-kb | del-primer_CEP128_440-kb_Fw | GCAGAATCATCCAGCGAATCAG |
| | del-primer_CEP128_440-kb_Rv | TGTTGGCATTCCTATCACGTC |
| CEP128 WT-primers 20-kb | WT-primer_CEP128_20_Fw | TTTCTTCGCATGAGTGGTGC |
| | WT-primer_CEP128_20_Rv | TGGCACTGGGTCAGGTAAAC |
| CEP128 WT-primers 50~440-kb | WT-primer_CEP128_50-440_Fw | CCACGTGACAAGAGAACCCA |
| | WT-primer_CEP128_50-440_Rv | TGAGATGCCCTTCGTGCAAT |
| HNRNPA1 del-primers 1000-kb | del-primer_HNRNPA1_1000-kb_Fw | CCCTAAACGTCCAACTGCCT |
| | del-primer_HNRNPA1_1000-kb_Rv | AACTGTCACCAACTGCCACA |
| MTAP_outside WT-primers | WT-primer_MTAP-outside_Fw | AGGTAGAGCCAGACTGGGAG |
| | WT-primer_MTAP-outside_Rv | TGCAGACCTTTCCGGTTCTC |
| MTAP WT-primers | WT-primer_MTAP_Fw | GGAGGCTTGTCTGCAGTAGG |
| | WT-primer_MTAP_Rv | AGCTTTACAAGTCCGGTGGG |
| CDKN2A WT-primers | WT-primer_CDKN2A_Fw | AGAATTCTCCCCCGTCCGTA |
| | WT-primer_CDKN2A_Rv | CGTTTCCTTCCTCCGCGATA |
| CDKN2B WT-primers | WT-primer_CDKN2B_Fw | CACTGAGGAGAGATTCCCGC |
| | WT-primer_CDKN2B_Rv | GACATCCCACGAGCCATCAT |
| DMRTA1 WT-primers | WT-primer_DMRTA1_Fw | AGACTTGACTGCGACCAAGG |
| | WT-primer_DMRTA1_Rv | GCATAATCCAAAGCTGGCCG |
| DMRTA1_outside WT-primers | WT-primer_DMRTA1-outsideFw | CCGAATCAAAGCATGGGCAC |
| | WT-primer_DMRTA1-outsideRv | ACACCCGAATCCCTAAGCAA |
| 9p21.3 del-primers | 9p21.3_del-primerFw | AGAGGGTACGCTTGCAAATGA |
| | 9p21.3_del-primerRv | AAAACACGTGGCTCGCCTA |

| Primer for knock-in check | Name | Sequence |
|---|---|---|
| HNRNPA1 Knock-in check primers | KI-primer_HNRNPA1_Fw | CAGGCCTTCAGCCGTTACAC |
| | KI-primer_HNRNPA1_Rv | GGGAGAATTACATGGGGAAGTGA |
